# Sex chromosomes, sex ratios and sex gaps in longevity in plants

**DOI:** 10.1101/2021.10.03.462936

**Authors:** Gabriel AB Marais, J-F. Lemaitre

## Abstract

In animals, males and females can display markedly different longevity (also called sex gap in longevity, SGLs). Recent work has revealed that sex chromosomes contribute to establishing these SGLs. X-hemizygosity and toxicity of the Y chromosomes are two mechanisms that have been suggested to reduce male longevity (Z-hemizygosity and W toxicity in females in ZW systems). In plants, SGLs are known to exist but the role of sex chromosomes remains to be established. Here, by using adult sex ratio as a proxy for measuring SGLs, we explored the relationship between sex chromosome and SGLs across 43 plant species. Based on the knowledge recently accumulated in animals, we specifically asked whether: (i) species with XY systems tend to have female-biased sex ratios (reduced male longevity) and species with ZW ones tend to have male-biased sex ratios (reduced female longevity), and (ii) this patterns was stronger in heteromorphic systems compared to homomorphic ones. Our results tend to support these predictions although we lack statistical power because of a small number of ZW systems and the absence of any heteromorphic ZW system in the dataset. We discuss the implications of these findings, which we hope will stimulate further research on sex-differences in lifespan and ageing across plants.

## 1. Introduction

In many animal species, males and females have different mortality patterns and longevities. This phenomenon has been observed in mammals, birds and in invertebrates (reviewed in (1)). For example, mammalian females generally outlive males, although some exceptions have been documented (2). Studying sex-differences in lifespan (also called sex gaps in longevity, SGL) is important for deciphering the physiological and genetic determinants of the aging process and also for developing sex-specific anti-aging interventions in humans (3, 4). Three non-exclusive possible evolutionary hypotheses underlying SGL have been proposed (see (1, 5)): the sex-specific long-term costs of sexual selection, the mother’s curse and the effects of sex chromosomes. In species where sexual selection strongly acts on males, the evolution of marked sexual dimorphism, sex-specific physiologies (e.g. high androgen levels in males) and conspicuous behaviours (e.g. sexual display, aggressiveness) can affect age-specific mortality patterns in a sex-specific way and lead to a reduced lifespan for males (6, 7);. Evidence accumulated so far have revealed that the level of sexual selection can - to some extent - explain the degree of SGL (6, 8), although this relationship can be largely modulated by environmental conditions (2). The mother’s curse relies on the asymmetric transmission of the mitochondrial genome (through females). Selection will thus be completely blind to all the deleterious mutations that will affect males only, which will favour accumulation of such mutations in the mitochondrial genomes and could potentially be detrimental for male’s survival prospect (9). However, while most empirical investigations of the mother’s curse have revealed detrimental effects of the accumulation of mtDNA mutations for male’s health and fertility (e.g. (10, 11)), impacts in terms of reduced male’s lifespan remain poorly documented (but see (12)). In theory, sex chromosome content could also affect aging and longevity (13). In highly heteromorphic systems, the Y chromosome is highly degenerated and most genes have an X copy only. Males are thus X-hemyzygous for most of the genes and they will thus express all the deleterious mutations (the so-called unguarded X effect, (13)). In females, recessive mutations on one X will often be masked by the other X. Moreover, the Y itself can have a toxic effect and reduce male lifespan (reviewed in (1)). Y chromosomes often includes large portion of DNA repeats including transposable elements (TEs). These elements can jump from one place to another in the genome causing somatic mutations and are often silenced via epigenetic mechanisms. The epigenetic marks can fade with time and in aging males, TEs can resume activity causing mutation rate acceleration, which can possibly lead to a faster aging and a reduced lifespan (1). While recent research suggests that all mechanisms are probably at work (1, 4), their relative contributions and how they may vary from one species to another is yet to be quantified.

The impact of sex chromosomes on SGLs is a new area of research. Before 2015, this mechanism was considered purely theoretical and some authors doubted that it even existed (13) (5). In 2015, Pipoly and colleagues documented a compelling association between adult sex ratios (used as a proxy for SGLs) and sex chromosome types (XY or ZW) in tetrapods (14). They found that the adult sex ratio was biased toward the homogametic sex (XX females or ZZ males) in amphibians, snakes and lizards, mammals and birds suggesting a higher morality for the heterogametic sex (XY males or ZW females) in agreement with both unguarded X and toxic Y mechanisms. This observation was later confirmed by a study using various metrics of longevity and conducted on a wider taxonomic range (i.e. vertebrate and invertebrate species, (15)). From these studies, however, it is not clear whether the unguarded X or the toxic Y effects are at work, or if both are what is their relative contribution. In *Drosophila melanogaster*, females outlive males and several studies have shown that the unguarded X has limited or no effect (16, 17) (but see (18)). The toxic Y is probably the main cause of the SGLs in *D. melanogaster* (19). TE activity increases faster in aging males compared to aging females and this correlates with chromosome compactness changes (loss of epigenetic marks typical of heterochromatin). Moreover, longevity is correlated with sex chromosome content not phenotypic sex: longevity is reduced in female XXY flies and is increased in male XO files (compared to female XX and male XY files respectively, (19)). The toxic Y effect has been confirmed in *Drosophila miranda* (20). A correlational study suggests that the toxic Y effect may be widespread also in tetrapods (21). They have indeed observed (but using a relatively small dataset, ca. 20 species) that Y chromosome size (relative to genome size) is correlated to male longevity in agreement with the toxic Y (but not expected under the unguarded X). This raises the possibility of an interplay between sex chromosome size and SGLs. Selection for male viability and longevity may favour shortening of the Y chromosome, an interesting possibility given that the fact that old and degenerated Y (or W) chromosomes are tiny remains a mystery.

On the other hand, SGLs have been poorly studied in plants. We know that plants can exhibit extreme differences in longevities among species, with herbs being annual plants and some trees being several hundreds or even thousands years old (e.g. (22–24)). Most flowering plants are hermaphrodites but 6% are dioecious, representing about 15000 species (25). A study of adult sex ratios on 243 of these plants has revealed that they likely exhibit SGLs (26). Growth forms (herb, shrub, tree, etc…) is one of the major determinants of the variation in SGLs among plants, with herbs tending to exhibit balanced sex ratios while trees exhibit strongly male-biased sex ratios, probably because female trees suffer from a substantial survival cost of reproduction. Interestingly, sex chromosomes were found to be associated with adult sex ratios. Species with documented sex chromosomes had a female-biased sex ratio, with the strongest biases in species with heteromorphic sex chromosomes. Whether XY and ZW systems show different correlation with sex ratios (as observed in animals, (14, 15) is currently unknown. The study of SGLs in dioecious plants is important because a global view on aging and its evolution ought to include plants. Moreover, plants are interesting because dioecy is derived contrary to the situation in animals (where the ancestor was gonochoristic and hermaphroditism is derived). Dioecious plants are thus good systems to study emerging SGLs. Also, sexual dimorphism is weak in plants (27), which make them potentially good systems to study the other mechanisms underlying SGLs. Of note, the mother curse may be strong as both mitochondria and plastids are transmitted maternally in plants. Finally, dioecious plants are over-represented among crops (17%, (28)), understanding and managing SGLs may be useful for agronomical purposes.

In the Field et al.’s dataset, information on sex chromosomes was available for only 22 species. No information was available on sex chromosome type (XY and ZW), precluding any analysis using this information. Since 2013, a number of studies have reported and characterized sex chromosomes in dioecious plants (see reviews from (29–31)). We thus updated the Field et al. (2013)’s dataset with this information. Our goal was to conduct an analysis on the sex ratios contrasting XY and ZW systems and to test the following predictions: under the toxic Y or unguarded X effects, the heterogametic sex should have a reduced lifespan, XY systems should show female-biased sex ratios, ZW ones should show male-biased sex ratios. These biases are supposed to be stronger in heteromorphic systems compared to homomorphic ones.

## 2. Methods

### 2.a. Dataset

We extracted adult sex ratio values from Field et al. (2013) (dataset available in DRYAD (https://datadryad.org/stash/dataset/doi:10.5061/dryad.28kd1). Adult sex ratios were computed as the number of males (individuals with male flowers) divided by the total number of individuals (males and females). Only populations with >10 sampled individuals were conserved for computing the adult sex ratios as in (26). We assumed that adult sex ratios were good proxies for SGLs. In other words, a female-biased mortality should lead to an adult sex ratio biased towards males (i.e. ASR > 0.5) while a male-biased mortality should lead to an adult sex ratio biased towards females (i.e. ASR < 0.5). We updated the information on sex chromosomes for all species included in the dataset. In particular, we sought for information about presence of sex chromosomes and sex chromosome type (XY/ZW and homomorphic/heteromorphic). We screened all papers using cytology/genetics/genomics (entered as keywords) published after 2012 on sex chromosomes and sex determination for the 243 species included in the Field et al. (2013)’s dataset using Pubmed and Google scholar. We also looked for species for which both sex ratios and information about sex chromosomes were published since 2012 but found only one species (*Amborella trichopoda*), which was added to the dataset. All data used in our analysis are provided in supplementary information.

### 2.b. Statistical procedure

To avoid any statistical issue due to phylogenetic inertia (32), we used Phylogenetic Generalized Least-Squares models (PGLS). The strength of the phylogenetic signal on the error structure of each model was assessed with the Pagel’s λ (33). Pagel’s λ was estimated with maximum likelihood by using the PGLS command from the R-package *caper* (34) and was further incorporated into the models to control for the phylogenetic dependence among species. The λ classically varies between 0 (no phylogenetic signal) and 1 (the observed pattern is predicted by the phylogeny; similarity among species scales proportionally to their shared evolutionary time following a Brownian motion model; see (33)). For all the phylogenetically-controlled analyses, data were linked to the phylogenetic tree built by (26). Yet, we modified this phylogenetic tree by pruning out the species without sex chromosomes and by adding *Amborella trichopoda* at the basis of the tree (*A. trichopoda* is known to be the sister species of all other extent angiosperms, (35, 36)) and obtained the tree shown in Figure 1). It is important to note that for all the models described below, λ was not statistically different from 0 and analyses performed without accounting for phylogenetic dependence among species lead to qualitatively similar results (data not shown).

**Figure 1:**
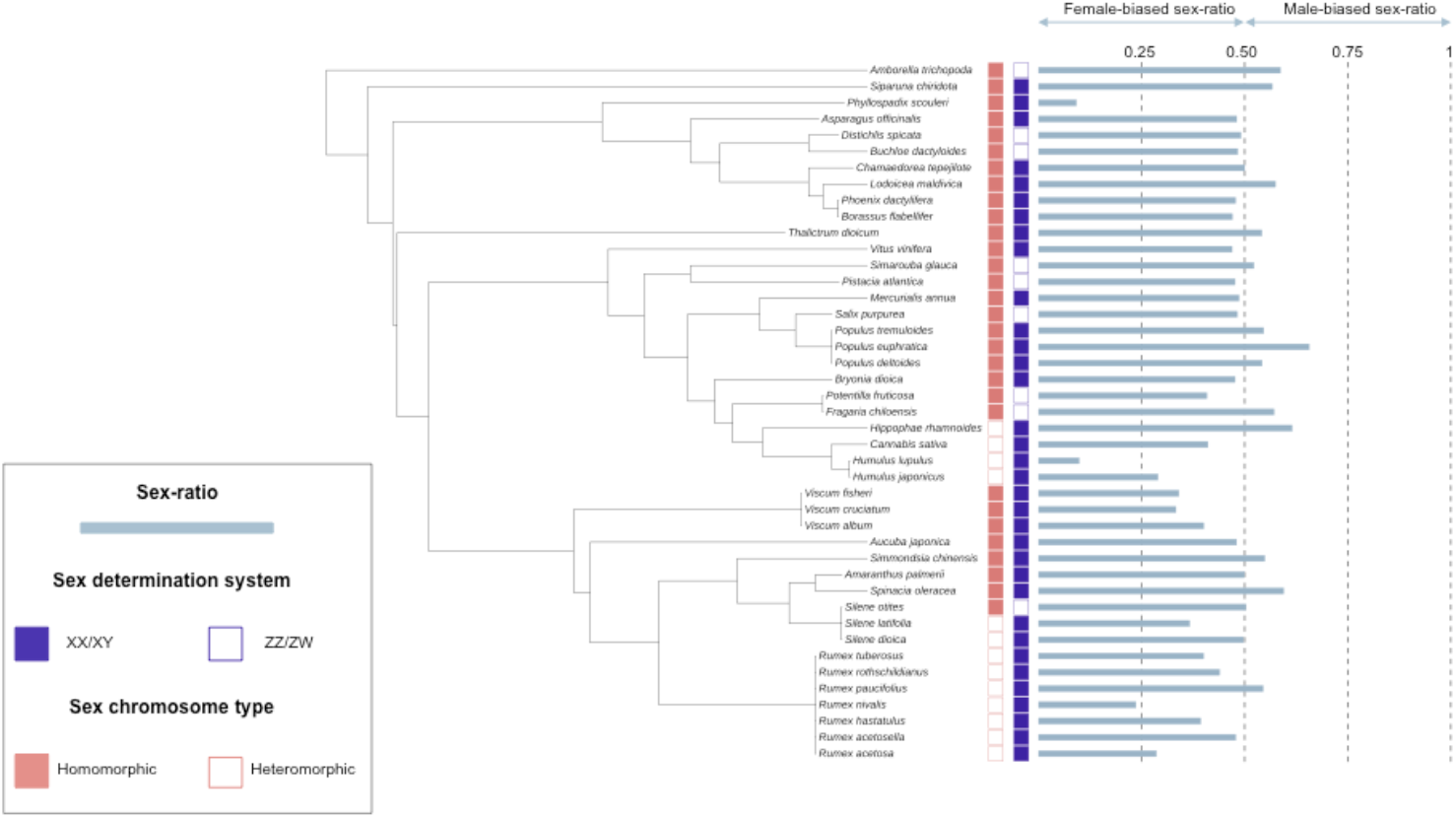
Phylogenetic tree of the species with sex chromosomes included in the dataset and used for the PGLS analysis.

To explore the association between ASR and sex chromosomes, we first analysed whether the variation in adult sex ratios observed among plants was explained by the genetic sex-determination system by fitting PGLS models with ASR as the response variable in all models, and the sex chromosome type (XY/ZW) as a predictor. Then, we tested whether the variation in adult sex ratios observed among plants differed between heteromorphic and homomorphic systems by fitting PGLS models with ASR as the response variable in all models, and the level of heteromorphy (heteromorphic/homomorphic) as a predictor. No heteromorphic ZW system was present in our dataset and we thus compared ASR values between the three following group of plants: homomorphic XY, heteromorphic XY and homomorphic ZW. All PGLS models were performed with R 4.0.0 (R Core Development Team) using the packages *ape* (37) and *caper* (34). Unless otherwise stated, parameter estimates are given as ± 95% Confidence Interval.

## 3. Results

Table 1 shows a summary of our work to update the Field et al. (2013)’s dataset with information on sex chromosomes. We doubled the number of species for which this information is available and the dataset now includes information on sex chromosome type. Our first analysis revealed that ASR did not differ between XX/XY and ZZ/ZW plants (Table 2A, Figure 2). However, we observed that species with XY systems have a slightly biased female-biased sex ratio (mean ASR = 0.44, 95% CI [0.40;0.49]; median ASR = 0.48, *N* = 34), while those with a ZW system have a balanced sex ratio (mean ASR = 0.5, 95% CI [0.46;0.54]; median ASR = 0.49, *N* = 9). Similar to what was observed with the whole dataset, ASR did not differ between XX/XY and ZZ/ZW in herb-like species (Table 2A, Figure 2). In herb-like species, species with XY systems have a slightly biased female-biased sex ratio (XY: mean ASR = 0.42, 95% CI [0.36;0.49], median ASR = 0.48, *N* = 17) while ASR did not deviate from 0.5 in ZW species (mean ASR = 0.51, 95% CI [0.44;0.57]; median ASR = 0.50, *N* = 4). However, in trees ASR confidence interval includes 0.5 in both groups (XY: mean ASR = 0.47, 95% CI [0.39;54]; median ASR = 0.48, *N* = 17; ZW: mean ASR = 0.49, 95% CI [0.41;0.58]; median ASR = 0.48, *N* = 5) highlighting an absence of deviation from a balanced sex ratio.

**Table 1:** Summary of the updating of the Field et al. (2013)’s dataset on dioecious plant sex ratios

|  | **Total** | **Sex**<br>**chromosomes** | **XY** | **ZW** |
| --- | --- | --- | --- | --- |
| **Field et al. (2013)** | 243 | 22 | n/a | n/a |
| **Updated Field et al. (2013)** | 244* | 43 | 34 | 9 |

\* Added species = *Amborella trichopoda* (Sex ratio from (38), sex determination type from (41))

**Table 2:**
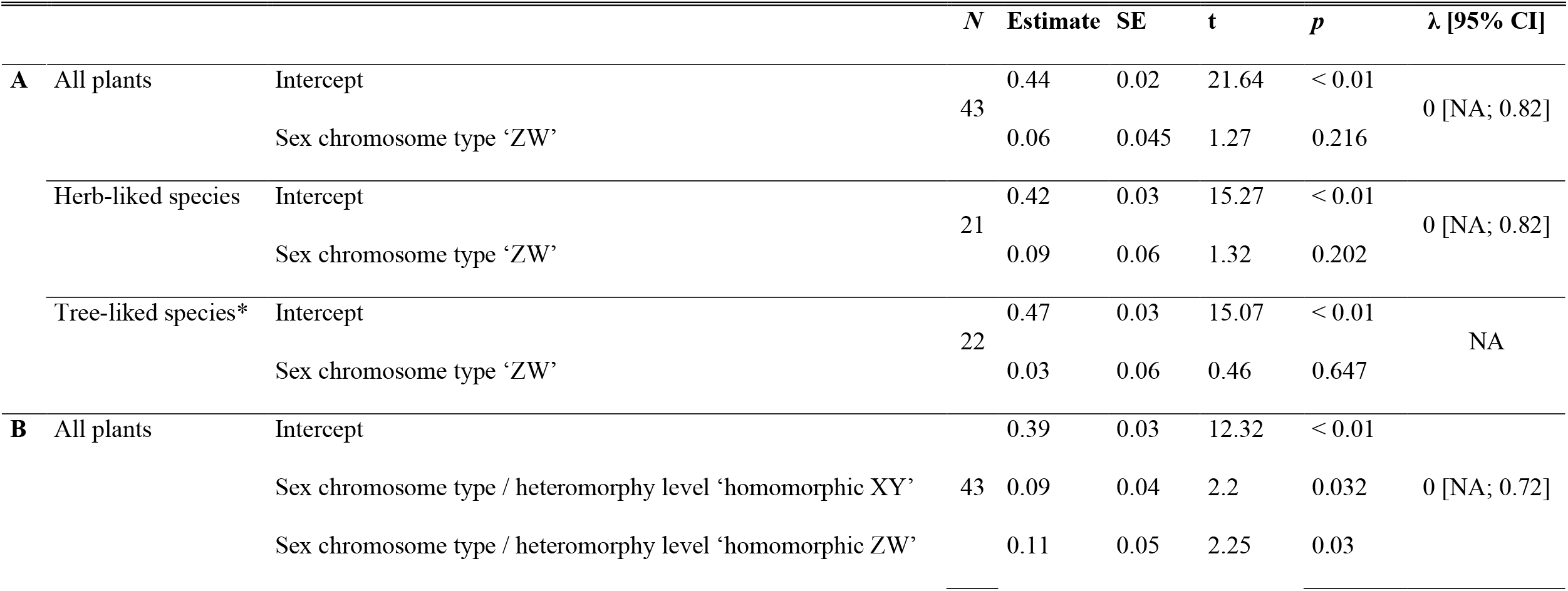
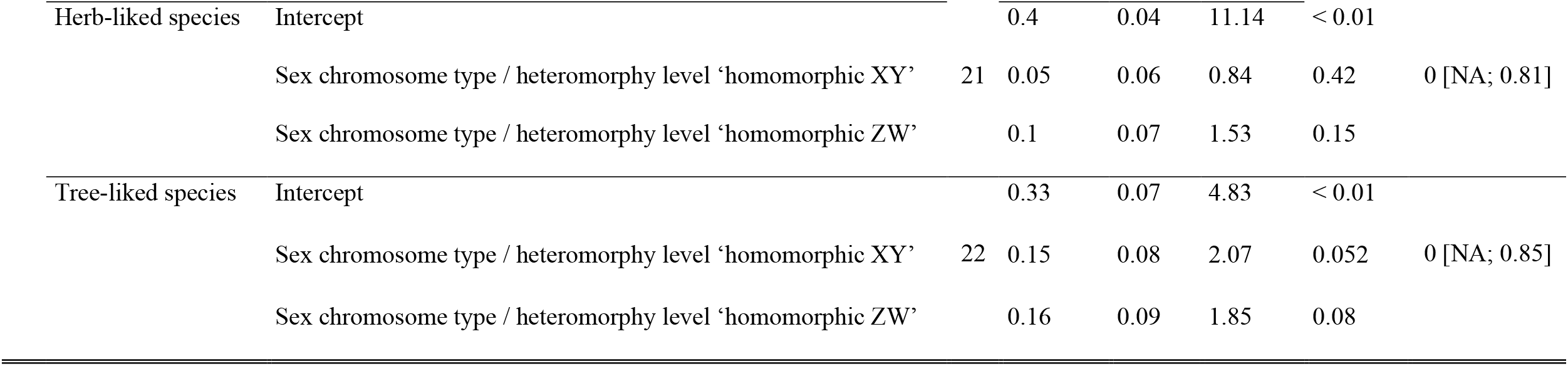
Parameters of the models discussed in the text. In all models the λ value was not statistically different from 0 indicating that the phylogeny did not influence the relationship between sex chromosomes and sex ratio. * indicates that the estimate corresponds to an ordinary least regression, instead of PGLS (due to lack of model convergence). In A, the sex chromosome type ‘XY’ has been used as a reference and in B, the sex chromosome type / heteromorphy level ‘heteromorphic XY’ has been used as a reference.

**Figure 2:**
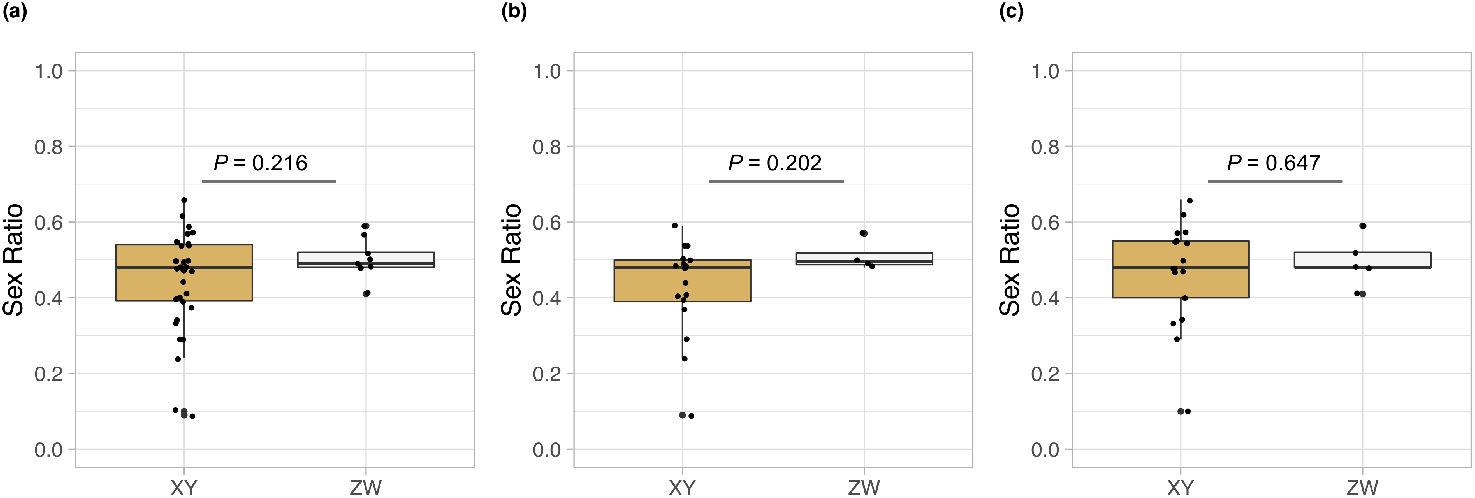
Sex ratios and sex chromosomes. A) all species, B) Herb-like plants, C) tree-like plants. Herb-like plants include annual and perennial herbs and tree-like plants include shrubs, trees, vines and mistletoes according to (26). P-values are indicated.

We then analysed separately homomorphic and heteromorphic systems. In the heteromorphic XY systems, we found a strongly female-biased sex ratio (mean ASR = 0.39, 95% CI [0.30;0.47]; median ASR = 0.40, *N* = 13). We found the same for herbs (mean ASR = 0.40, 95% CI [0.34;0.47]; median ASR = 0.41, *N* = 10) while in trees the sample size was too low to draw any definitive conclusion, despite a female-biased sex ratio (mean ASR = 0.33, 95% CI [-0.31;0.99]; median ASR = 0.29, *N* = 3). Homomorphic systems on the other hand exhibit close-to-balanced sex ratios (mean ASR = 0.49, 95% CI [0.45;0.52]; median ASR = 0.49, *N* = 30; see Figure 3). As expected, our analyses revealed that the level of heteromorphy influence ASR (Figure 3). ASR values were significantly lower in heteromorphic XY than in homomorphic sex chromosomes (Table 2B). When analyzed separately, a similar pattern was found in tree-liked plants and in herb-liked plants, although it was less clear in the latter (Table 2B).

**Figure 3:**
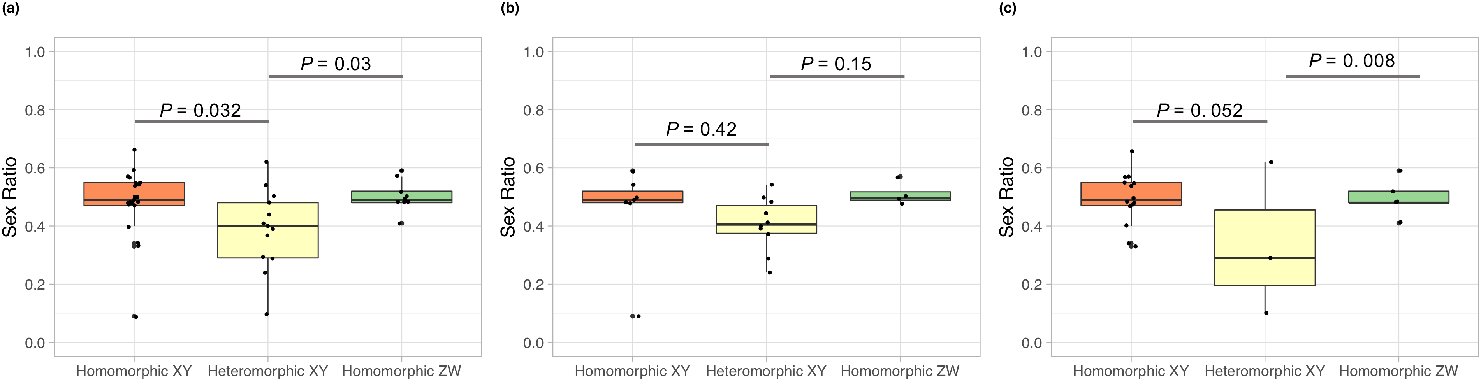
Sex ratios and sex chromosome heteromorphy. A) all species, B) Herb-like plants, C) tree-like plants. Herb-like plants include annual and perennial herbs and tree-like plants include shrubs, trees, vines and mistletoes according to (26). P-values are indicated.

## 4. Discussion

Our results support an effect of sex chromosomes in contributing to establish SGLs in plants. The heterozygous sex seems to experience reduced lifespan. This is clear for XY systems and less clear for ZW ones in our results, possibly due to lower representation of the latter in our dataset. The patterns are stronger for heteromorphic systems, although this could be observed for XY systems only, as our dataset only included homomorphic ZW systems. There is, to our knowledge, no documented heteromorphic ZW system in plants yet. Patterns look similar in herbs and trees, and are perhaps more pronounced in trees. This was somewhat surprising as Field et al. (2013) found that overall sex ratio biases have different direction in herbs and trees: female-biased in the former, male-biased in the latter (26). We observe that sex ratios were more female-biased in trees and herbs with heterogametic XY systems, suggesting that such sex chromosomes have a strong impact on male survival and possibly aging in trees.

Of course our dataset only included 43 species, which limits its statistical power. Moreover, adult sex ratios are very rough proxies for SGLs. They are correlated to a number of factors, perhaps the main one being growth form, which we took into account here. But there are also affected by sex ratios at germination, and adult sex ratios are valuable proxies for SGLs only if sex ratio at germination is 1:1. Biased sex ratios at germination are known to exist in plants (discussed in (26)), although there are also reports of unbiased ratios in many plants (e.g. (38, 39)). Future studies should ideally use larger datasets (including more ZW systems) and other proxies (demographic data).

Our study aims at stimulating more research on the effect of sex chromosomes on sex-specific differences in longevity in plants. Sex chromosomes are being characterized in a growing number of plant species (29–31), which should help updating the Field et al.’s dataset further and also enrich it with information on the size of the non-recombining region, the level of Y degeneration, the age of the sex chromosome system. Getting more sex ratios data (or longevity data) would also be useful. These are lacking for some interesting species. *Coccinia grandis* for example is the dioecious plant with the strongest documented sex chromosome heteromorphy (40), and a strong SGL is expected for this plant but no sex ratio data are currently available in this species.

## Ethics

N/A

## Data accessibility

All data used for the analysis performed in this manuscript are provided in supplementary information.

## Authors’ contributions

Conceptualization: GAB; Data curation: GABM; Formal analysis: JFL; Funding acquisition: GABM, JFL; Investigation: GABM, JFL; Visualization: JFL, GABM; Writing – original draft: GABM; Writing – review & editing: GABM, JFL

## Acknowledgements

We thank Editors Susanne Renner and Niels Müller for their positive feedback about this work and the possibility to contribute to this special issue. We thank all the members of the LongevitY ANR grant for discussions.

